# Mapping uterine calcium dynamics during the ovulatory cycle in live mice

**DOI:** 10.1101/2024.02.02.578395

**Authors:** David J. Combs, Eric M. Moult, Sarah K. England, Adam E. Cohen

**Affiliations:** Department of Anesthesiology, Perioperative and Pain Medicine, Brigham and Women’s Hospital, Harvard Medical School; Department of Chemistry and Chemical Biology, Harvard University; Department of Obstetrics and Gynecology, Center for Reproductive Health Sciences, Washington University School of Medicine; Department of Physics, Harvard University

## Abstract

Uterine contraction patterns vary during the ovulatory cycle and throughout pregnancy but prior measurements have produced limited and conflicting information on these patterns. We combined a virally delivered genetically encoded calcium reporter (GCaMP8m) and ultra-widefield imaging in live nonpregnant mice to characterize uterine calcium dynamics at organ scale throughout the estrous cycle. Prior to ovulation (proestrus and estrus) uterine excitations primarily initiated in a region near the oviduct, but after ovulation (metestrus and diestrus), excitations initiated at loci homogeneously distributed throughout the organ. The frequency of excitation events was lowest in proestrus and estrus, higher in metestrus and highest in diestrus. These results establish a platform for mapping uterine activity, and show that the question of whether there is an anatomically localized trigger for uterine excitations depends on the estrous cycle phase.

## Introduction

Uterine contractions support important biological functions. In pregnancy, they are central to labor and delivery, as well as postpartum hemostasis (1). Outside of pregnancy, uterine contractions assist in fertilization, implantation and the clearance of menstrual debris (2). Aberrant uterine contractility is implicated in preterm birth, dystocia, postpartum hemorrhage, infertility, dysmenorrhea, and endometriosis, all common conditions, many of which carry significant morbidity or mortality.

Cellular and molecular studies have identified many of the ion channels and other proteins central to uterine myocyte electrophysiology (3). However, it is not clear how these components produce tissue-scale activity, and how that activity is coordinated and controlled during ovulation and pregnancy. As contractions in pregnancy are quasi-periodic (4) and non-pregnancy contractions vary in frequency and origin with the ovulatory cycle (5–11), prior work sought evidence for uterine pacemakers (12). However, without high-quality organ-scale maps of uterine activity patterns, the search for a putative uterine pacemaker is difficult. Past searches have largely assumed similarity to known pacing systems (*e*.*g*. heart or gut) and looked for analogous electrical activity or histologic features (13, 14), yielding conflicting views. High resolution activity maps in live animals could pinpoint the loci driving organ-wide activity and could inform future searches for the molecular mechanisms underlying the cycle-dependent changes in activity patterns.

Measurement of calcium dynamics with genetically encoded calcium indicators presents a solution to this challenge. These tools are readily deployed at organ scale in living animals and have been used in brain (15), mammary ducts (16), skeletal muscle (17), and heart (18, 19). In combination with cell type-specific drivers, these indicators can report on the activity of cellular subpopulations within a complex tissue.

Here, we use the genetically encoded calcium indicator jGCaMP8m to map myometrial calcium dynamics across an entire uterine horn in live mice at different stages of the estrous cycle. These measurements clarified the nature of cycle-dependent changes in propagation velocity, locus of initiation, and the event rate. We also identified a dominant locus of initiation for excitation events in the ovarian end of the uterine horn, but it was aperiodic and active only before and during ovulation (proestrus and estrus) and not afterward (metestrus and diestrus).

## Methods

### Mouse strains and maintenance

All animal procedures adhered to the National Institutes of Health Guide for the care and use of laboratory animals and were approved by the Harvard University Institutional Animal Care and Use Committee. Progesterone Receptor (PR)-Cre^+/-^ mice (JAX 017915) were generously provided by the strain’s creator, Nirao Shah at Stanford University (20). Anti-Müllerian Hormone Receptor 2 (AMHR2)-Cre+/-mice (21), were generously provided by the laboratory of Kara McKinley at Harvard University. Ai148 mice (JAX 030328), for cre-dependent GCaMP6f expression, were obtained from the Jackson Laboratory. All mice were housed in standard conditions (reverse 12-hour light/dark cycles, with water and food *ad libitum)*. For all experiments involving injection of viruses carrying *cre-*dependent payloads, genotyped female heterozygotes were generated from crosses with wild-type C57BL/6 mice obtained from Charles River Laboratories. Genotyped PR-Cre^+/-^;Ai148D^+/-^ or AMHR2-Cre^+/-^;Ai148D^+/-^ double heterozygotes were created by the crossing of respective parental strains. All genotyping was performed by Transnetyx.

### Vaginal cytology and estrous cycle staging

Vaginal cytology and estrous staging were performed by vaginal lavage with 50-100 μL of phosphate buffered saline (PBS) using a micropipette tip and suction bulb followed by crystal violet staining of freshly dried cytology preparations, as previously described (22). All cytology specimens were collected approximately 1 hour before lights-off. For virus-injected female mice, first cytology specimens were not obtained until at least 21 days post-operatively to allow for surgical recovery and restoration of uninterrupted light-dark cycling. Estrous staging was determined by daily vaginal cytology over at least five days prior to measurement. Calcium imaging of a particular estrous stage was only performed when the day-of-measurement cytology matched the cytology expected based on the prior days’ specimens (see Supplemental Materials).

### Virus injection surgery

AAV9 virus expressing jGCaMP8m under a CAG promoter was obtained from Addgene (162381-AAV9). For injections, pre-aliquoted frozen virus was thawed and diluted to 5x10^12^ GC/mL in a total volume of 10 μL, containing 10% (v/v) of trypan-blue dye to visualize spread of the injectate and to guide adjustments to the injection location(s).

For the virus injection procedure (see also Supplemental Materials), 8-12 week old heterozygous PR-Cre^+/-^ female mice were anesthetized, a vertical incision was made left paramedian to the midline to access the peritoneal cavity and the left uterine horn identified. Injections were made using home-fabricated micropipettes (Sutter P1000) with beveled tips created with a home-built micropipette polisher. Micropipettes were mounted in a microinjection pump (WPI Nanoliter 2010), positioned with a micromanipulator (WPI, MP-285), and injection volumes controlled with a microsyringe pump controller (WPI, Micro4). After loading of viral injectate and positioning of the micropipette near the uterine horn, injections were performed by gentle grasping of the mesometrium with micro forceps and simultaneous advancement of the micropipette into the uterine horn. Virus was injected in two 300 nL volumes at 50 nL/sec at each injection site with 5-10 seconds of post injection time before pipette removal to assess injectate spread and prevent leakage. Injection sites consisted of 12-16 evenly spaced locations along the anterior face of the organ longitudinal axis from the left half of the uterine body (junction of the two uterine horns) to the horn end adjacent to the oviduct. Following injections, the fascial layer was closed with interrupted 5-0 monocryl sutures and the skin closed with wound clips.

### Animal preparation for live animal imaging

For live animal imaging, after isoflurane anesthesia (see Supplemental Material), a midline abdominal incision was made to expose the peritoneal cavity and the bladder drained transmurally. A pair of custom 3D-printed (Anycubic, Photon Mono) semi-circular (16 mm diameter, 8 mm height) abdominal retractors were positioned at the inferior and superior borders of the abdominal incision, with extension of the incision to accommodate retractor insertion (∼3.5 cm). Soaked gauze strips were adjusted to sit between the retractor and the abdominal wall to protect the wall surface from drying or damage from the retractors. The time from induction of anesthesia to collection of the last image was tracked for each animal and kept under 90 minutes. It was generally not possible to image the entirety of both uterine horns simultaneously, so for animals where both the injected and contralateral horn were imaged, gauze and retractors were adjusted to optimize the view of the imaged horn. After each animal imaging session, the animal was euthanized.

### Calcium imaging

The *in vivo* imaging stage was positioned below the objective of a custom optical setup for wide-field imaging previously described (23, 24). For recordings of jGCaMP8m dynamics, widefield illumination was supplied (0.1-0.2 mW mm^-2^) with a 470 nm LED. Excitation light was reflected off a large (65 × 80 × 3 mm) dual band (509-540 nm, 614-700 nm, Alluxa) dichroic mirror. Fluorescence was separated from back-scattered excitation by a 65 mm diameter 520/35 nm band-pass emission filter. Imaging was done via a 1X objective lens, numerical aperture 0.25 (Olympus MVPLAPO 1X). Data were captured on a Kinetix sCMOS Camera (Teledyne Photometrics) at 5 Hz framerate through a camera lens repurposed as a tube lens (VS Technology Co, VS-50085/M42). The iris was set to achieve the focal depth needed to image the convex organ surface and to accommodate any anteroposterior displacement along the organ longitudinal axis. The adjoining oviduct, junction of the two uterine horns and (often 1-2 mm) of the contralateral horn could also be imaged simultaneously. Illumination and acquisition timing was controlled using custom MATLAB software (https://www.luminosmicroscopy.com) via a National Instruments DAQ interface. For details on image processing see Supplemental Materials.

### Immunostaining and confocal microscopy

For immunostaining of uterine rings (see Supplemental Materials), rings were permeabilized (0.1% Triton X-100 in PBS), blocked (1% bovine serum albumin in PBS with 0.1% Tween 20), and then incubated with the primary antibody in fresh blocking solution for 1 hr at room temperature. After secondary antibody incubation (1 hr, room temperature), rings were incubated for 5 minutes with 300 nM DAPI, washed, and mounted on glass slides. Primary antibodies were: chicken anti-GFP (Abcam, ab13970, 1:500) and rabbit anti-alpha smooth muscle actin (Abcam, ab124964, 1:500). Secondary antibodies were donkey anti-chicken Alexa Fluor 488 (Jackson Immunology 703-545-155, 1:1000) and anti-rabbit Alexa Fluor 647 (Abcam, ab150075, 1:1000).

Imaging of fixed immunostained uterine rings was performed on an LSM 700 Confocal Microscope and Zen v3.2 software (Zeiss). Image stacks were converted to maximal intensity projection images in ImageJ.

### Statistics

Plots were created in MATLAB. All boxplots show median and interquartile range (IQR) with whiskers indicating 1.5 x IQR. The numbers of independent repeat experiments performed in different animals are displayed as *n* values. For statistical comparisons of features between estrous stages that proved significantly different (p < 0.05) by 1-way ANOVA, Tukey’s test was used with a multiple comparison procedure. For paired comparison of the injected and contralateral horn, two tailed paired t-testing was performed. Sample size was determined by the practical limitations of the experiments. Researchers were not blinded.

## Results

### Adeno-associated virus injection enables in vivo myometrial transgene expression

To map uterine calcium dynamics, we developed methods for *in vivo* transgene delivery and calcium imaging. Adeno-associated virus (AAV)-mediated transgene expression has not been reported in uterine smooth muscle. Plasmid-based uterine muscle-specific promoters have not been described, and no transgenic mice have been created that express site-specific recombinases exclusively in uterine smooth muscle. We therefore chose a commercially available AAV with payload comprising the universal CAG promoter and Cre-dependent expression of the calcium indicator jGCaMP8m (25). We chose the AAV9 capsid serotype, which has been successfully used for *in vivo* transgene expression in skeletal and cardiac muscle (26). We used progesterone-receptor (PR)-Cre mice to target expression to myometrium (20). Virus was injected into the anterior face of the uterine horn along its longitudinal axis via an open abdominal surgery (Figure 1; Methods). In virus-injected animals, estrous staging was determined by vaginal cytology (Figure 1, Supplemental Figure 1; Methods).

**Figure 1:**
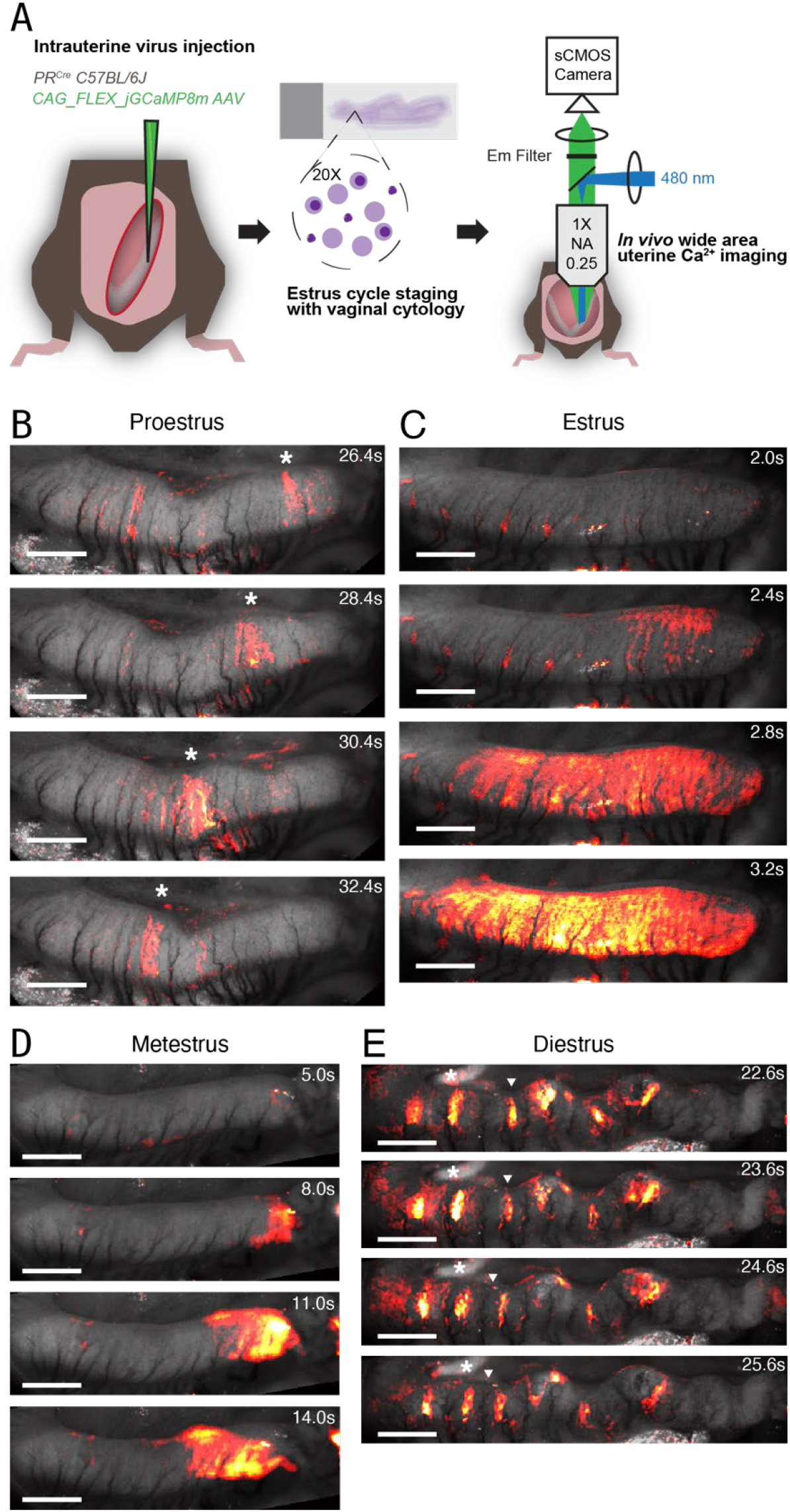
Live animal wide-area uterine calcium imaging during estrous cycle. (A) Experimental scheme. Left panel: The uterus was injected with AAV virus encoding GCaMP8m. Middle: Estrous stage was determined with vaginal cytology. Right: Uterine calcium dynamics were imaged on an ultra-widefield microscope. Images from movies of calcium dynamics with the change in fluorescence in color overlaid on a grayscale image of the basal fluorescence for (B) proestrus, (C) estrus, (D) metestrus, and (E) diestrus. Asterisks (B and E) and arrowheads (E) mark propagating bands of activity in successive images. Scale bars 1 mm in all images.

Virus injection led to expression of jGCaMP8m in the injected, but not the contralateral, uterine horn (Supplemental Figure 2). Despite injections only in the anterior face of the uterine horn, jGCaMP8m expression spread around the circumference of the organ. jGCaMP8m expressed in both circular and longitudinal muscle layers (Supplemental Figure 2). Consistent with the known tissue-specific expression pattern of Cre in PR-Cre mice (27), jGCaMP8m also expressed in the endometrium. Expression lasted for at least 12 weeks after injection. Injected uteri appeared anatomically normal (Supplemental Figure 3). Virus injection and transgene expression did not produce detectable adverse impacts on reproduction, as pregnancy in virus-injected animals was readily achieved (n = 9) in the same time frame (< 3 months post injection) as our imaging experiments.

We also tried crossing two Cre driver lines (PR-Cre and AMHR2-Cre) with a Cre-dependent GCaMP6s reporter mouse line to drive whole-uterus reporter expression. Female offspring expressed calcium indicator throughout their uteri, but these animals had atrophic uteri and were infertile, so this strategy was not pursued further (Supplemental Figure 4 and Supplemental Materials). Attempts to deliver transgenes by direct electroporation into wild-type uteruses were also not successful.

In addition to live animal imaging (see below), we tried imaging whole-uterus explants, but despite an extensive search of conditions, we found that the explanted uteri always showed markedly less spontaneous activity than *in vivo*. Mechanical or electrical stimuli could evoke activity in the explants, confirming that the tissue was alive (Supplemental Figure 5 and Supplemental Materials).

### Wide-area live animal calcium imaging reveals estrous-related changes in calcium dynamics

We imaged uterine calcium dynamics *in vivo* in mice expressing virally delivered jGCaMP8m (Figure 1B-E, Supplemental Movies 1-4). Fluorescence dynamics in the injected horn were much larger than the small motion-associated changes in tissue autofluorescence from the contralateral horn (Supplemental Figure 6). We observed different kinds of calcium excitation events. Some excitations comprised elevated calcium in rings around the tissue, coupled with a local reduction in organ diameter (Figure 1B, 1D). Other excitations increased calcium in axially elongated strips, coupled with long-axis shortening (Figure 1C,1E). We provisionally associate these events to independent activation of the circular and longitudinal muscle layers, respectively.

Recordings of calcium dynamics were taken from mice in all four estrous stages (*n* = 5 mice per stage). Qualitatively, videos from proestrus and estrus mice had markedly less activity than those from metestrus and diestrus mice (Supplemental Movies 1-4). To analyze differences in activity among the four estrous stages, we developed an image processing pipeline to extract and quantify uterine excitation events (Supplemental Figure 7, Supplemental Materials). In brief, we used motion-tracking and non-rigid deformation algorithms to undo the effects of tissue motion. We then defined a piecewise linear contour along the axis of the expressing uterine horn, and projected the fluorescence dynamics onto this axis, leading to fluorescence kymographs comprising one time dimension and one spatial dimension along the uterine axis (Figure 2A). This procedure discarded information about circumferential calcium dynamics, but visual inspection of the movies showed that most of the propagation was axial. We then used a combination of machine learning-based (28) and feature-based algorithms to identify calcium events and to quantify their locus of initiation, propagation speed, propagation duration, full-width at half maximum, (Figure 2, Figure 3, Methods).

**Figure 2:**
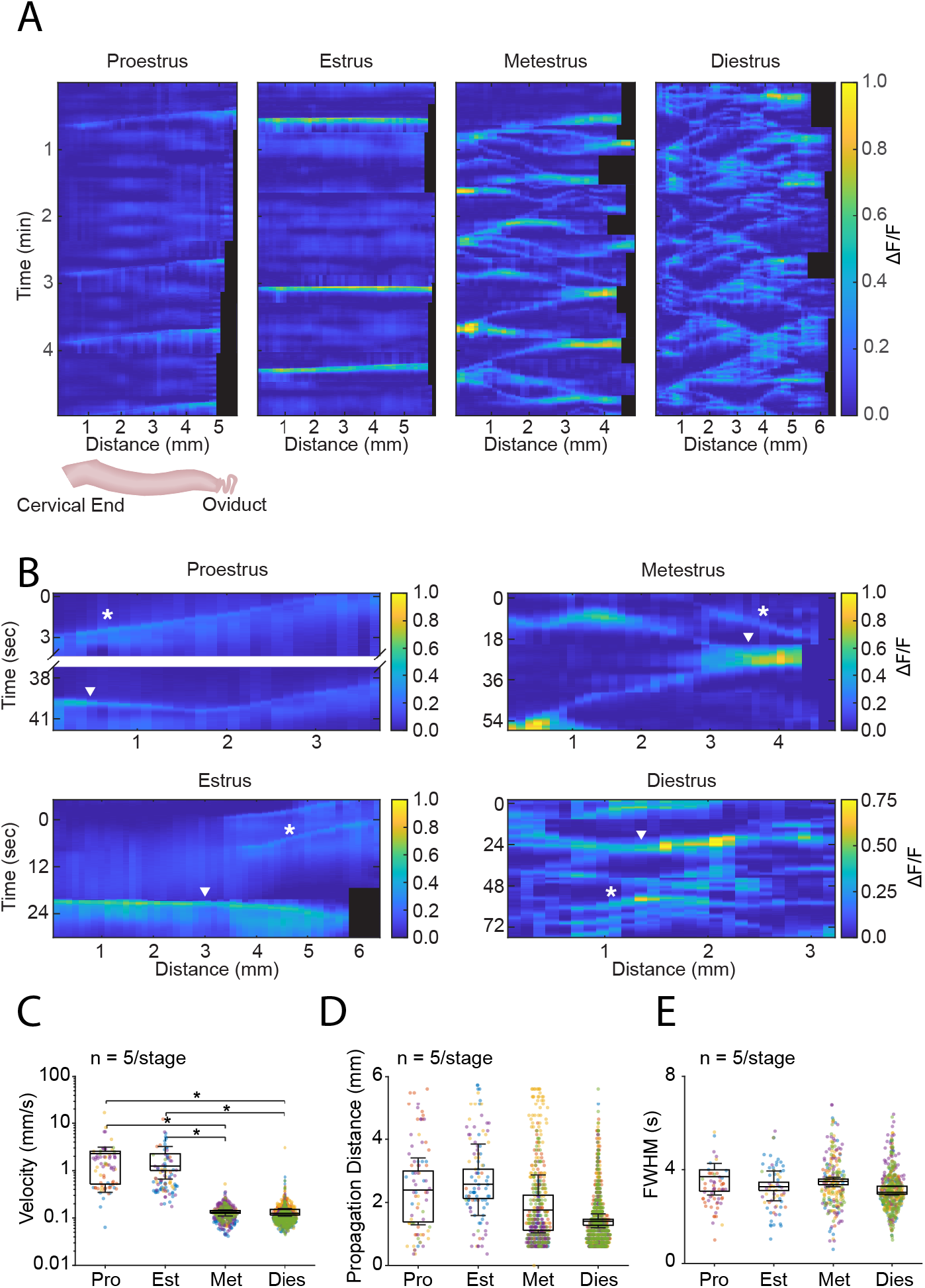
Uterine calcium dynamics vary by estrous stage. (A) Representative kymographs of fold-change in fluorescence for each estrous stage. (B) Kymograph segments showing fast (arrowhead) and slow (asterisk) conduction modes in same specimen. For each panel in A and B, the right-most spatial coordinate is closest to the oviduct. Black regions indicate organ length contractions, leading to no data from the ovarian end. Event-wise distributions of (C) propagation velocity, (D) distance and (E) full width at half maximum (FWHM). Box plots represent IQR (25^th^, 50^th^, and 75^th^ percentile) with whiskers representing 1.5 IQR. Swarm plot points represent values from individual calcium events. Colors identify events occurring in the same animal. Proestrus (Pro), estrus (Est), metestrus (Met) and diestrus (Dies) stages had n = 5 animals per stage. For statistics (B), P < 0.01 (1 way ANOVA), with multiple-comparisons testing by Tukey’s test where *P < 0.05.

**Figure 3:**
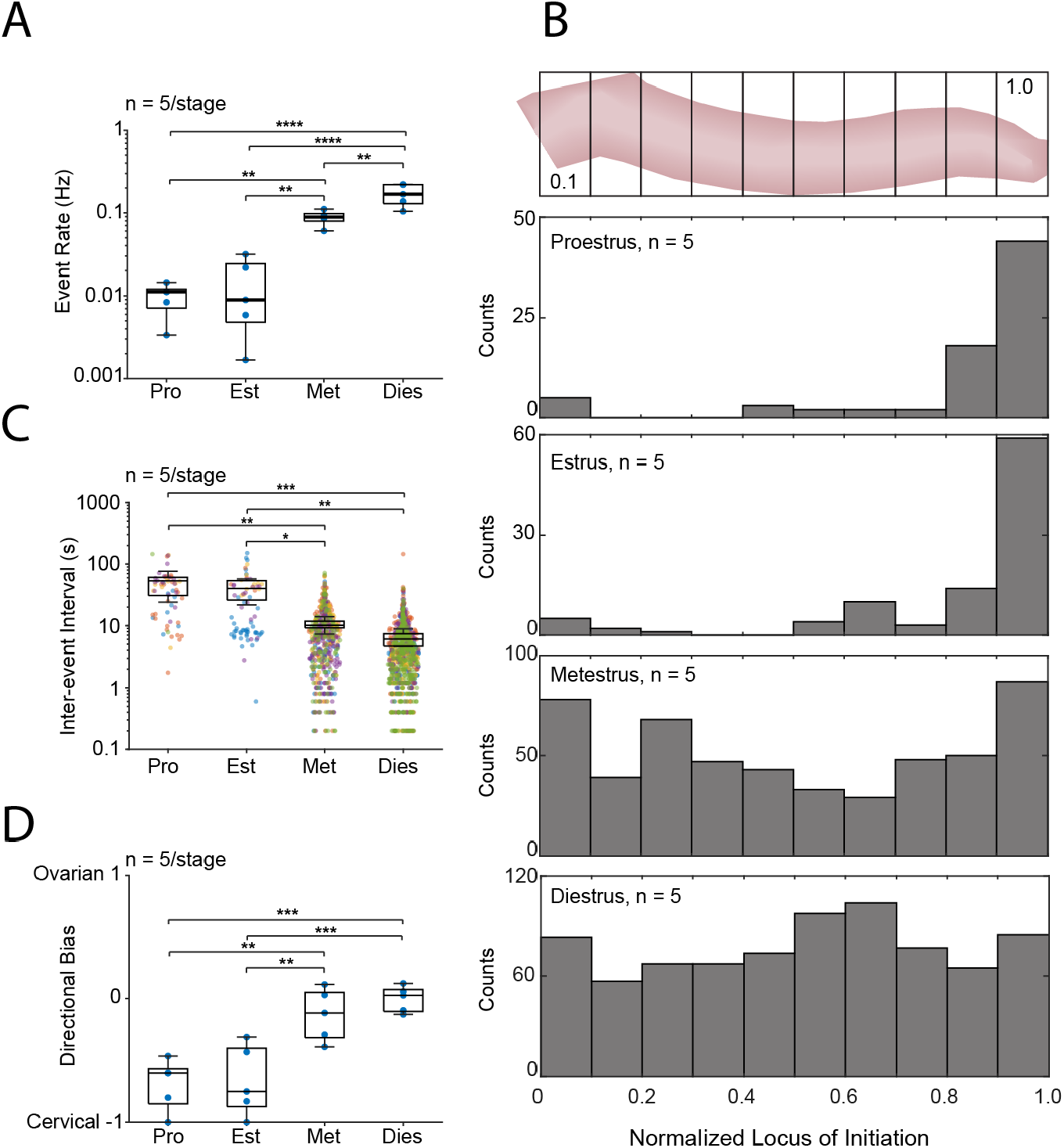
Frequency, direction, and loci of initiation of uterine calcium events vary by estrous stage. Distributions of (A) event rate, (B) inter-event interval throughout the estrous cycle. (C) Histograms of event loci of initiation, normalized relative to uterine horn length and binned into deciles as indicated in the top panel. Counts represent all events across 5 animals per stage. (D) Distributions of propagation directional bias. In (A) and (D), data points represent individual animals; in (C) data points represent pairs of successive calcium events. Like-colored points in (B) identify events occurring in the same animal. Box plots represent IQR (25^th^, 50^th^, and 75^th^ percentile) with whiskers representing 1.5 IQR. Proestrus (Pro), estrus (Est), metestrus (Met) and diestrus (Dies) stages had n = 5 animals per stage. For statistics, (A) P < 1 10^-6^, (C) P < 0.002, and (D) P < 1 10^-4^ by 1 way ANOVA; multiple-comparisons testing by Tukey’s test where *P < 0.05, **P < 0.01, ***P < 0.001, ****P < 1 10^-4^.

Individual kymographs revealed co-existence of quickly propagating and slowly propagating conduction modes, a feature observed in all cycle stages (Figure 2B). Examination of raw calcium recordings and organ motion suggested that these modes reflected activation of the longitudinal and circular muscle layers respectively. Since assignment of excitation events to these categories was sometimes ambiguous, subsequent analysis did not distinguish event types.

We compared the properties of calcium events across estrous stages. Propagation velocities in proestrus (2.3 ± 1.1 mm/s, median ± s.d.) and estrus (1.2 ± 0.9 mm/s) were faster than in metestrus (0.13 ± 0.01 mm/s) and diestrus (0.12 ± 0.02 mm/s; p < 0.01, 1 way ANOVA, Figure 2C). Median event propagation distances (as magnitudes) in proestrus and estrus appeared slightly higher than in metestrus and diestrus, but this did not achieve statistical significance (Figure 2D). To look at the duration of events at a given location (*i*.*e*. at a particular kymograph spatial coordinate), event-triggered averages were generated from their leading half maxima. Full widths at half maximum (FWHM), a measure of the duration of calcium elevation at a given location, of these mean event traces did not differ by estrous stage (Figure 2E). Event rates (number of events/total recording time, Figure 3A) were much lower for proestrus (0.01 ± 0.004 Hz) and estrus (0.009 ± 0.01 Hz) than for metestrus (0.09 ± 0.02 Hz) and diestrus (0.17 ± 0.05 Hz; p < 1x10^-6^). Table 1 summarizes these observations.

**Table 1:** Velocities and event rates of calcium events.

| **Cycle Stage** | **Velocity (mm/s)** | **Event rate (Hz)** | **Origin** |
| --- | --- | --- | --- |
| **Proestrus** | 2.3 ± 1.1<sup>a</sup> | 0.01 ± 0.004 | Near oviduct |
| **Estrus** | 1.2 ± 0.9 | 0.01 ± 0.01 | Near oviduct |
| **Metestrus** | 0.13 ± 0.01 | 0.09 ± 0.02 | Distributed |
| **Diestrus** | 0.12 ± 0.02 | 0.17 ± 0.05 | Distributed |

a. Median and standard deviations

### A region with pacer-like features exists in proestrus and estrus, but not metestrus and diestrus

In the heart, an anatomically and molecularly distinct pacemaker region coordinates the beating of the whole organ. Whether an analogous structure exists in the uterus has been controversial (4, 12). We searched for pacemaker-like dynamics in the uterine Ca^2+^ signal. Two key elements of a pacing system are localization and periodicity. That is, events may originate from a single locus or from multiple loci; and may occur periodically or at irregular intervals.

To examine where events initiated, we mapped our data onto a normalized uterine horn longitudinal axis, and then sorted event LOIs into deciles along this axis. During estrus and proestrus, activity primarily initiated from regions near the oviduct, with 82% and 74% of events, respectively, emerging from two deciles closest to the oviduct (Figure 3B). In contrast, events in metestrus and diestrus showed an even distribution of LOIs across the uterine horn (Figure 3B). The spatially distributed LOIs in diestrus and metestrus appeared equally likely to propagate toward the oviduct or toward the cervix, whereas those in proestrus and estrus showed a strong directional bias to propagate in the cervical direction (Figure 3C).

Although portions of some kymographs showed periodic activity emanating from a particular locus, this periodicity did not appear stable. To examine the temporal structure of events, we generated power spectra from kymographs of recordings at each estrous stage (Figure 4). The power spectra of dynamics in the uterine horn did not have any peaks, indicating absence of periodicity at any estrous stage. This observation was consistent with the wide distribution of inter-event-intervals for all four estrous stages (Figure 3D). To confirm that we could detect a periodic signal, we examined the oviduct, which, in a subset of samples, exhibited jGCaMP8m signal, likely acquired from viral injections into the oviductal end of the uterine horn. Consistent with previous reports of myosalpinx contractions in the oviduct paced by interstitial cells of Cajal (29), the oviductal calcium signal was highly periodic with a typical frequency of 0.15 Hz (Figure 4). We did not observe sustained synchrony of oviductal and uterine activity in any estrus stage.

**Figure 4:**
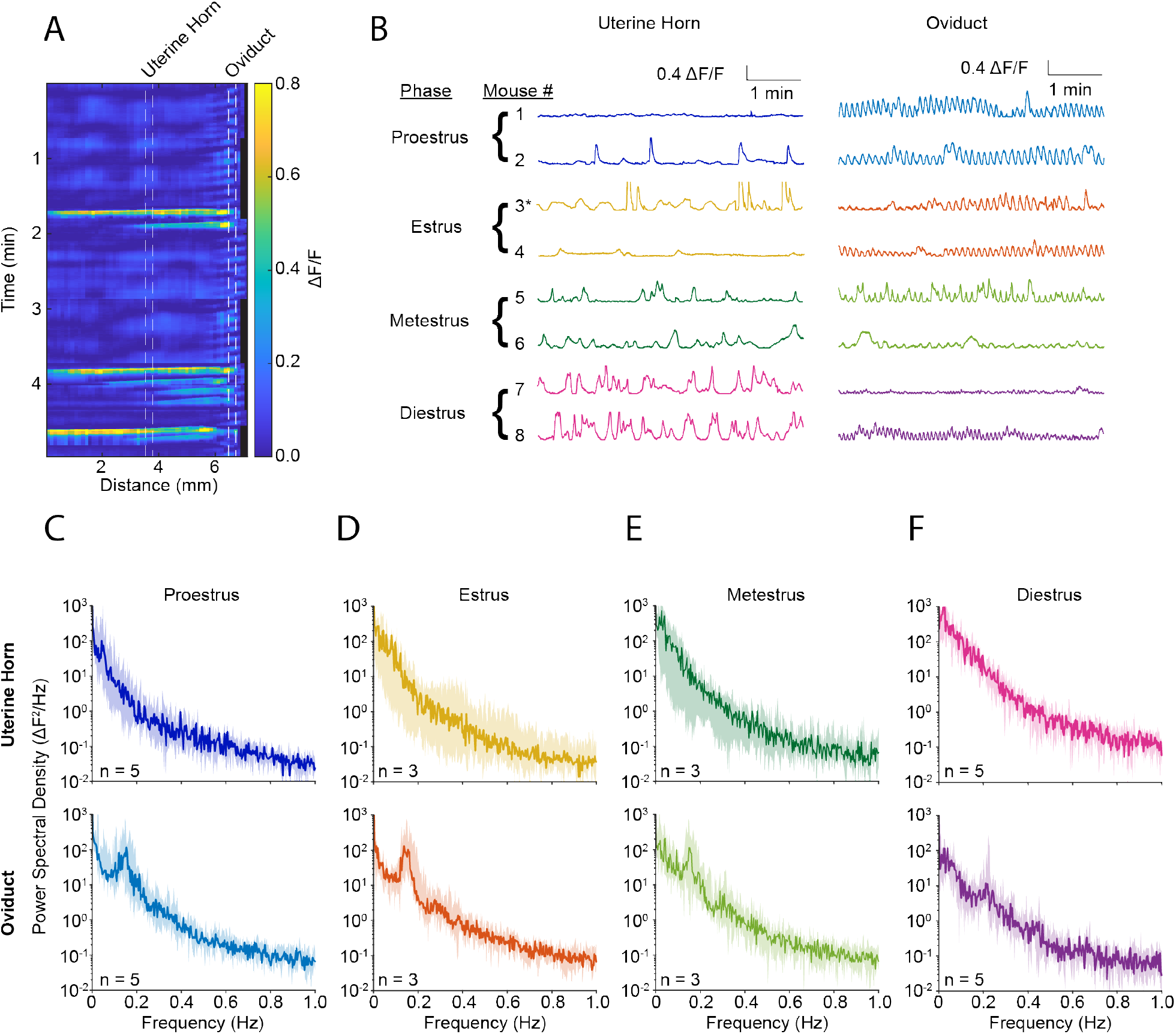
Uterine calcium dynamics are not periodic. (A) Kymograph with overlaid marks indicating regions analyzed for uterine horn and oviduct. (B) Representative fluorescence traces from two mice in each estrous stage. Asterisks indicate traces derived from the kymograph in (A). (C-F) Power spectral density of the uterine horn (top panels) and oviduct (lower panels) for each stage of estrous. The number of animals is marked on each panel.

## Discussion

We developed a protocol for expressing transgenes in the myometrium and for live-animal imaging of uterine calcium excitation dynamics across the organ. Uterine calcium activity rose following ovulation, peaking in diestrus. Proestrus and estrus stages featured rare, fast-propagating, and short-lived activity which started near the oviduct. Metestrus and diestrus exhibited more frequent, slower-moving, and longer-lasting excitations which initiated throughout the organ. These events propagated in equal amounts toward the cervix and toward the ovaries.

In the heart, a pacemaker drives excitations with a regular rhythm from a defined locus of initiation. In the uterus, we observed a preferred locus of initiation during proestrus and estrus, but not during other phases and the activity was never periodic. Thus the ovarian end of the uterus has some pacemaker-like properties some of the time, but does not contain a structure analogous to the anatomically and molecularly defined cardiac pacemaker. The physiological basis of the shift toward ovarian-end initiation during proestrus and estrus is not known. Single cell transcriptomic analysis of juvenile and adult mouse uteri identified four myocyte subtypes (30). Future studies should compare the gene expression patterns and electrophysiology of the ovarian end vs. the bulk horn, during epochs of localized vs. delocalized event initiation.

The high propagation velocities and lower event rates we observe in proestrus and estrus are consistent with previously reported increases in gap junction expression in those phases (31). Increases in gap junction conductance are expected to speed electrical conduction, but also to increase electrotonic loading of nascent initiation sites and thereby decrease the event rate (32). Differences in event rates may also be partially explained by the known and antagonistic effects of progesterone and 17β-estradiol on L-type calcium channel expression, leading to elevated calcium channel current densities in diestrus (33). Potassium channel subtypes with faster activation kinetics predominate in estrus vs. diestrus, which could shorten action potential and burst durations in estrus (34).

The circular and longitudinal layers of the rodent myometrium have different activity (35, 36), and we observed different conduction modes for organ level calcium dynamics within individual specimens. We did not attempt to sort calcium events by layer, but videos of dynamics suggested that the slow-propagating activity in metestrus and diestrus was mostly from the circular layer, while in proestrus and estrus, fast-propagating longitudinal layer events occurred at a similar rate as circular layer events. Cycle-dependent shifts in activity between these two muscle layers may also partially explain the changes we observed in overall activity patterns.

Prior studies reported seemingly inconsistent patterns of estrous cycle-dependent uterine dynamics in rodents. We propose that these differences are likely due to the use of different measurement modalities. Tissue motion, intra-uterine pressure, electrical recordings, and calcium imaging all probe different aspects of myometrial activity: different patterns of excitation could have stronger impact on some measures than others. Early electromyogram (EMG) studies in live rodents showed drops in activity from estrus to diestrus (5, 6). However, combining EMG with intrauterine pressure (IUP) measurements showed that not all electrical activity produced pressure changes (6). Furthermore, electrodes primarily report the activity of the cells they contact, and may be less sensitive or insensitive to activity in a different muscle layer. Microelectrode arrays provided in-layer spatial resolution and showed that estrus explants fired more synchronously than diestrus explants (11), consistent with the stage-dependent differences in event velocity we reported.

Our study confirms that of Wray and Noble, who studied tension and calcium in rodent myometrial strips and found diestrus to be the most active stage (9). In contrast, Dodds and coworkers used tension and motion measurements on uterine explants from mice and reported diestrus as the most quiescent phase (10). However, kymographs of uterine contraction from the Dodds study showed subtle chevron-like patterns matching our diestrus calcium kymographs. We infer that the highly localized circular-layer calcium events which we observed produced only small global changes in tension, pressure, and organ profile. Indeed, in our diestrus videos, slowly propagating circular-layer events only decreased uterus diameter by 0.1 mm ± 0.06 mm (median ± s.d.; *n* = 10 events, 5 animals), whereas circular-layer events in proestrus or estrus decreased organ radius by 0.4 mm ± 0.15 mm (*n* = 10 events, 6 animals). A recent study of organ motion in uterine explants also identified diestrus as the most active stage (37).

Our wide-area, live-animal calcium imaging provides a direct measure of myometrial excitation with high spatial and temporal resolution and addresses possible artifacts from prior measurement techniques. Prior experiments with dissected preparations cut across uterine muscle fibers (8–11), creating damage that may disrupt electrical activity. Even without dissection, explanted organs may be deprived of circulating factors or paracrine or neural stimuli. We observed that whole uterus explants did not reproduce native patterns of activity. Prior measurements of live rodent uterine activity (7, 38, 39) imposed substantial uterine stretch, likely altering endogenous dynamics (40). In our measurements, computational correction for tissue motion relieved the requirement to physically immobilize the tissue.

The biological significance of the distinct modes of uterine excitation we observed is not clear. While mice, like most mammals, do not menstruate, proliferating endometrial tissues degenerate and are ultimately resorbed in diestrus (41). We speculate that bidirectionally propagating activity in metestrus and diestrus may assist endometrial breakdown. Nonmechanical effects of calcium excitation events are also possible, e.g. via calcium-mediated changes in gene expression.

Unlike mice, women have a single fused uterine cavity without distinct circular and longitudinal muscle layers (2). However, analogously to the rodent estrous cycle, human myometrial activity varies over the menstrual cycle, with the strongest activity occurring at menstruation (42). The molecular mechanisms that augment myometrial activity following ovulation could be common to both species. However, perhaps because it is periodic, most studies of human non-pregnant myometrial activity have focused on the peristaltic activity of the thin layer of subendometrial muscle which is most active in estrus (43, 44). It is unclear whether mice have a similar layer, though mice may share similar hormonally sensitive activating mechanisms. In any case, our study emphasizes activity of bulk uterine muscle during endometrial decline, which in humans is connected to dysmenorrhea and endometriosis, common debilitating conditions.

Our study emphasizes the importance of using *in vivo* preparations for faithfully characterizing spontaneous activity patterns, and our platform has general applications for studying uterine physiology, pathophysiology, and pharmacology. In future studies, we plan to investigate the molecular mechanisms that underlie estrous-related changes and to adapt our approaches to studying the myometrium throughout pregnancy and in models of disease.

## Supporting information

Movie S1: Proestrus

Movie S2: Estrus

Movie S3: Metestrus

Movie S4: Diestrus

## Acknowledgements

We thank Shahinoor Begum and Andrew Preecha for technical assistance and mouse husbandry. We thank Kara McKinley and Nirao Shah for generously providing transgenic animals. We thank Ronald McCarthy for technical advice on mouse surgery and uterine tissue dissection. This work was supported by an award (D.J.C.) from Harvard Anesthesia Research Training Grant (T32, GM007592-45), the Young Investigator Award (D.J.C.) from the Society for Obstetric Anesthesiology and Perinatology, and an award (D.J.C.) from the Harvard BIRCWH Program (K12, AR084230-20).

