## Supplementary Materials for "Mapping uterine calcium dynamics during the ovulatory cycle in live mice"

Supplemental Video captions 1 – 4
Supplemental Figures 1 – 7
Supplemental Methods
Supplemental Discussion

**Supplemental Video 1: Live animal calcium dynamics during proestrus.** Scale bar 1 mm.

**Supplemental Video 2: Live animal calcium dynamics during estrus.** Scale bar 1 mm.

**Supplemental Video 3: Live animal calcium dynamics during metestrus.** Scale bar 1 mm.

**Supplemental Video 4: Live animal calcium dynamics during diestrus.** Scale bar 1 mm.

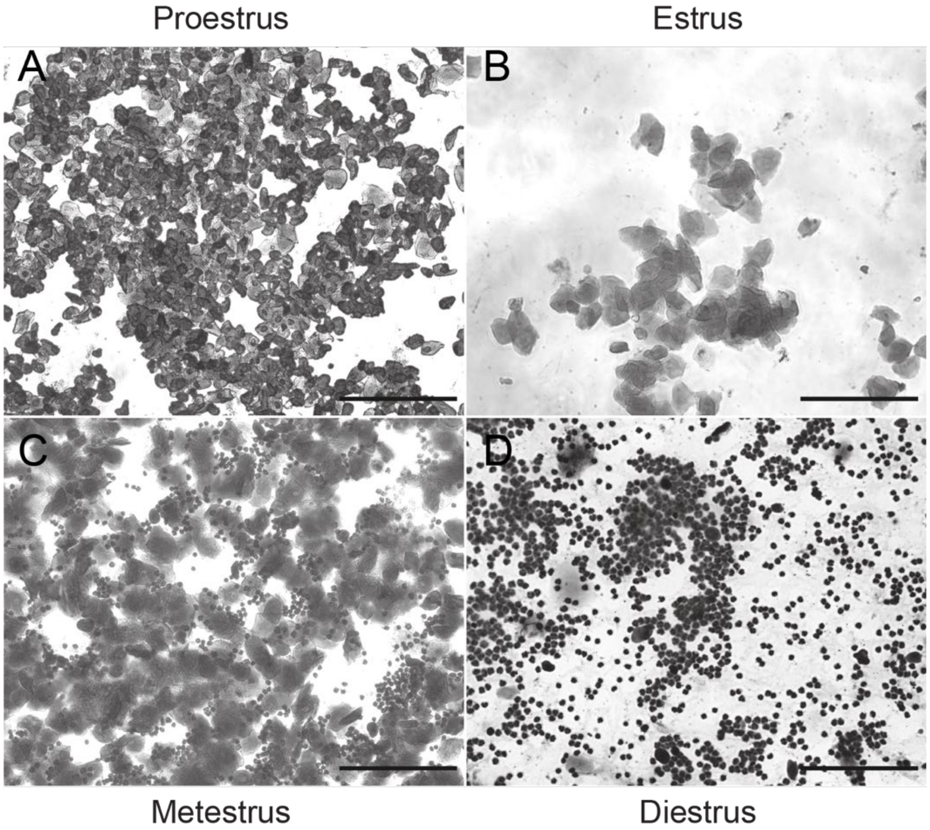


**Supplemental Figure 1: Vaginal cytology.** Representative brightfield images of vaginal cytology specimens for proestrus (A), estrus (B), metestrus (C), and diestrus (D). Scale bars 100 microns.

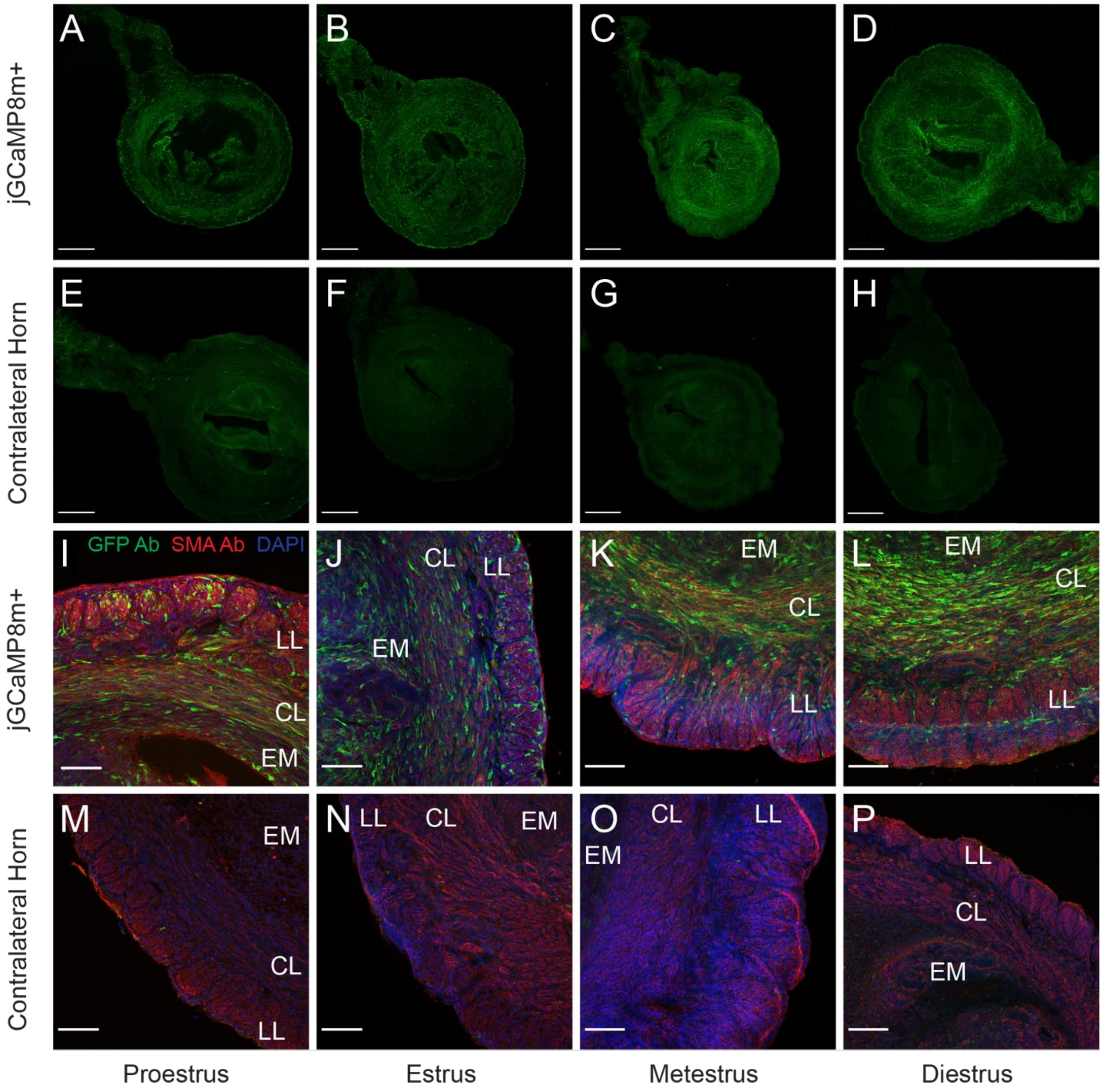


**Supplemental Figure 2: Confocal microscopy images of fixed sectioned rings of uterine tissue expressing GCaMP8m.** Images of tissue rings cut from (A-D) virus-injected and (E-H) contralateral non-injected uterine horns. Tissues were stained with anti-GFP antibodies and imaged with a 10X objective. Scale bars 500 microns. Maximum intensity projections of z-stacks of (I-L) virus-injected and (M-P) contralateral non-injected sections stained with anti-GFP and anti-SMA antibodies and DAPI and imaged with a 20X objective. Longitudinal muscle (LL), circular muscle (CL) and endometrial (EM) layers are labeled. Scale bars 100 microns.

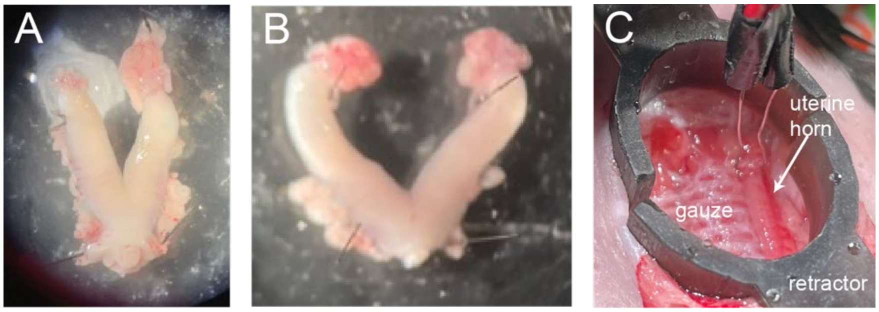


**Supplemental Figure 3: Photographs of explanted virus-injected uteri and live animal imaging preparation.** Injected (right) and contralateral (left) appear similar in diestrus (A) and estrus (B). Live animal imaging preparation (C) with black plastic retractors to expose uterus and underlying gauze to minimize drying. Home built stimulating electrode present near the oviduct was not used in this report.

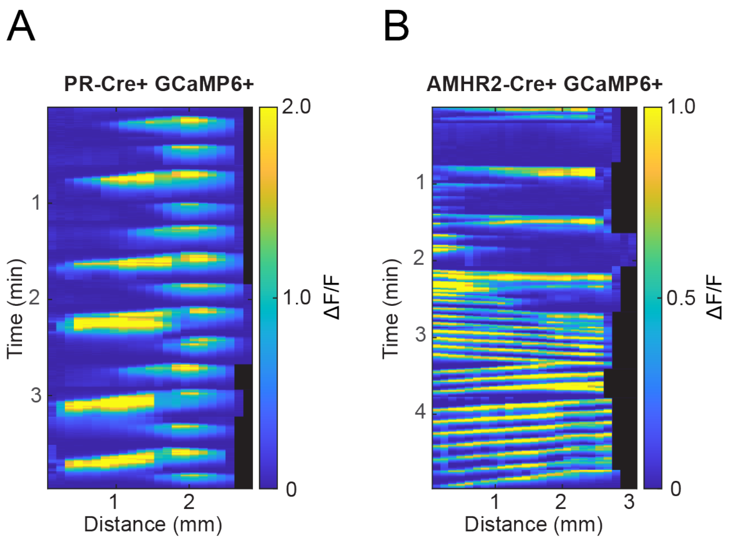


**Supplemental Figure 4: Kymographs from GCaMP6f-expressing transgenic animals.** Tissue-scale calcium signals were detectable in transgenic mice expressing Cre-dependent GCaMP6f driven by cells expressing (A) the progesterone receptor or (B) the anti-Müllerian hormone receptor 2. For each kymograph, the right-most spatial coordinate is closest to the oviduct. Black regions indicate organ shortening. Animals constitutively expressing transgenic GCaMP6f were not fertile and were not used for analysis.

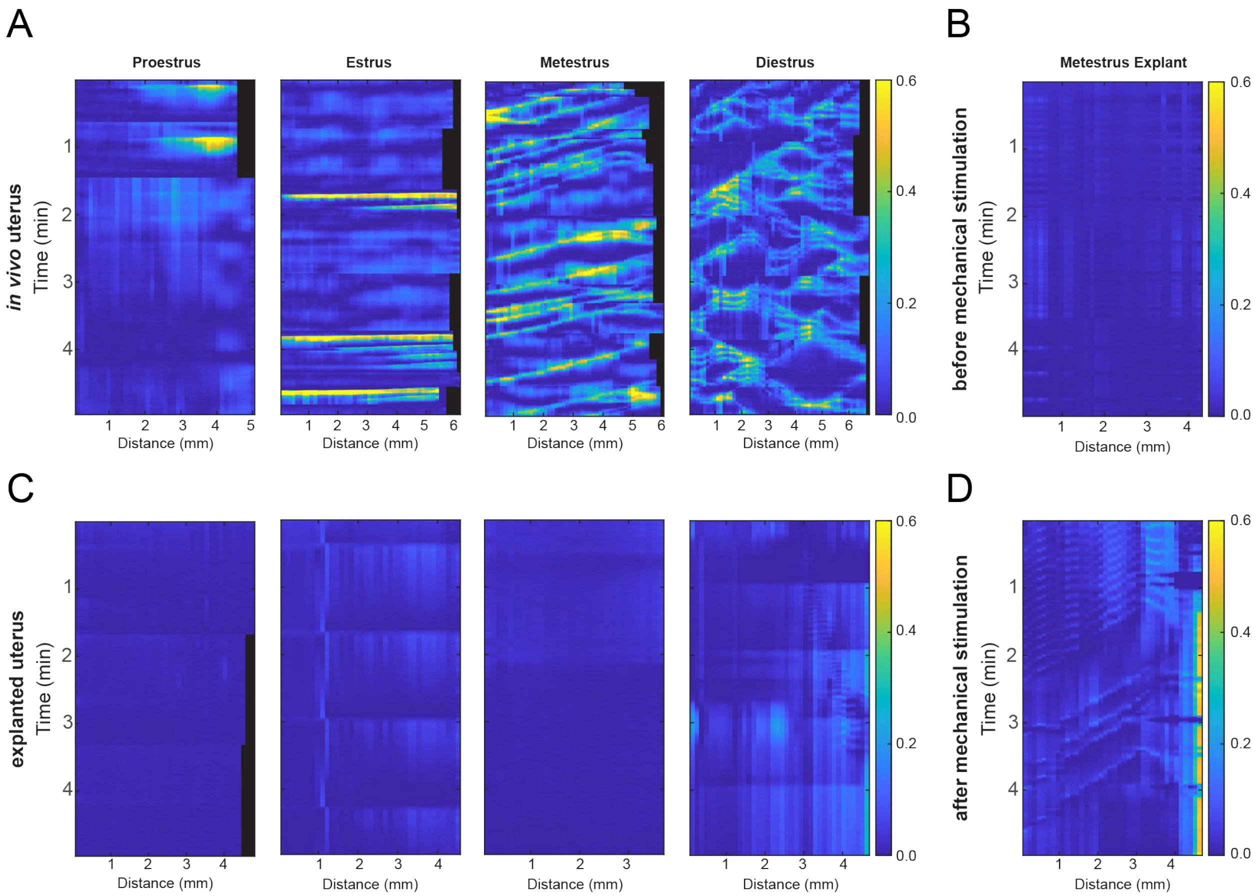


**Supplemental Figure 5: Uterine explants have markedly different dynamics than *in vivo*.** (A) Representative kymographs of *in vivo* uterine calcium dynamics for each estrous stage and (C) after explant. Note the faint endogenous calcium dynamics in diestrus stage explant (C, right-most panel) occurring between 2 and 4 minutes. (B) Representative kymograph of a uterine explant before and (D) after mechanical stimulation achieved by gentle tugging and bending of the imaged horn. For all kymographs, the right-most spatial coordinate is closest to the oviduct.

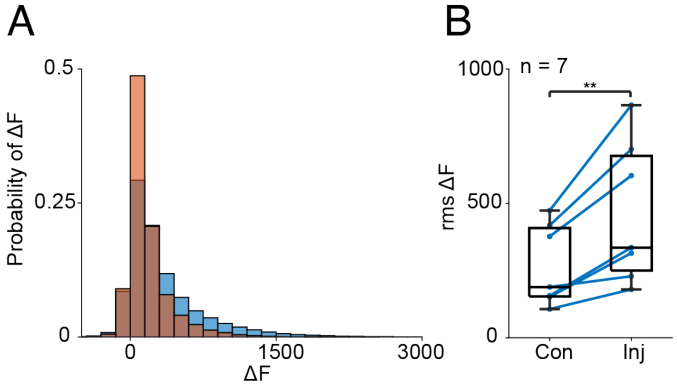


**Supplemental Figure 6: Fluorescence signals from injected horn exceed those of the contralateral horn.** (A) Histograms of the change in fluorescence for the injected (blue) and contralateral (red) horns. (B) Distributions of the root-mean-squared change in fluorescence for the injected (Inj) and contralateral (Con) horns where connected dots represent the two horns of individual animals (n = 7 animals). Box plots represent IQR ($25^{th}$, $50^{th}$, and $75^{th}$ percentile) with whiskers representing $1.5 \times$ IQR. Paired t test with **P = 0.005

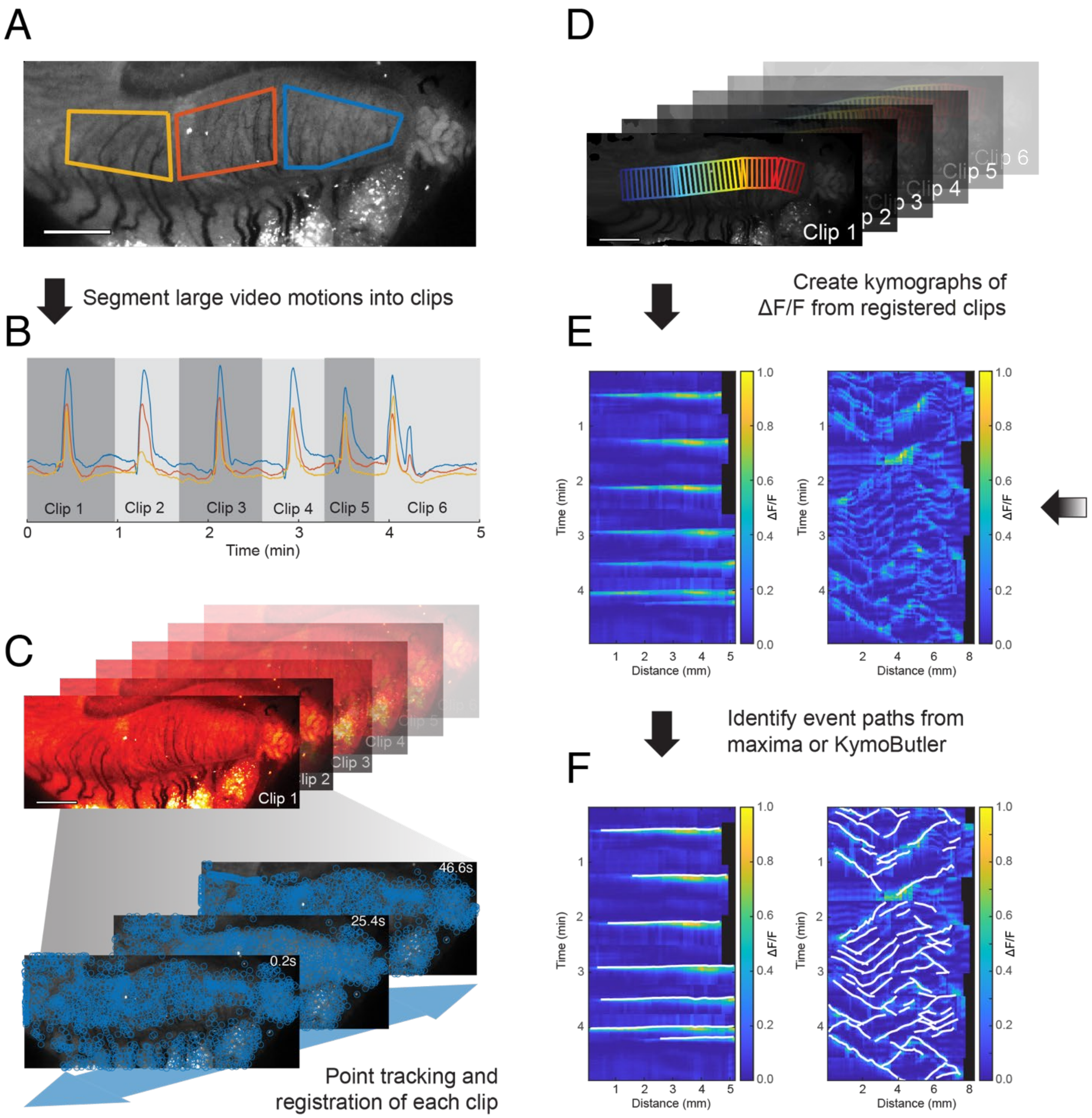


**Supplemental Figure 7: Scheme for analysis of calcium dynamics.** (A) Regions of interest (ROIs) were drawn to segment uterus into thirds. (B) Mean intensities from pixels in each ROI were plotted to identify episodes of motion and to segment videos into clips. (C) Anatomical features in each clip were marked with points which were tracked across successive frames to correct for uterine motion. (D) ROIs were tiled across motion-corrected clips, and the median intensity determined for each ROI. (E) The fold-change fluorescence of median intensity for each ROI in time was plotted as a kymograph. Black regions indicate organ shortening. (F) Event paths for individual kymographs were identified from event maxima for low activity proestrus and estrus uteri or the machine-learning tool KymoButler for high activity metestrus and diestrus uteri.

### Supplemental Methods

*Vaginal cytology*

Colony-housed female mice are known to cycle irregularly (1), so we adopted the following approach to improve the reliability of our stage determinations. To confirm the presence of cycling in a given female, cytology specimens were obtained no more than once daily over a minimum of five days until both an early stage (either proestrus or estrus) and a late stage (either metestrus or diestrus) were observed with the expected temporal separation, assuming a 4-5 day cycle (2). Females meeting this standard were deemed to be cycling. Then, to establish the estrous stage for an *in vivo* imaging experiment in a particular cycling female, vaginal cytology was followed no more than once daily over succeeding days. For a particular day, the stage was predicted based on the most recent day's result and the anticipated 4-5 day cycle. If that day's cytology result matched the predicted stage, it was accepted as the correct stage and the animal was used for *in vivo* imaging.

If the resulting cytology did not match the predicted stage, that mouse was not used for *in vivo* imaging that day. Instead, we continued to perform no more than daily cytology checks until we confirmed the start of a new cycle, we then attempted the prediction procedure again. In cases where experimental females were found not to cycle, to stop cycling, or to cycle irregularly, dirty male bedding was added to their cage to promote regular cycling (3), and our cytology approach was restarted.

*Approaches to genetically encoded reporter expression*

We also examined other methods of transgene expression. PR-Cre mice were crossed with a mouse line which expresses GCaMP6f under dual control of Cre recombinase and doxycycline (Ai148D; Methods). PR-Cre$^{+/-}$;Ai148D$^{+/-}$ double heterozygotes expressed GCaMP6f in uterine smooth muscle and showed clear uterine calcium dynamics (Supplementary Figure 5). However, these animals were undersized, died more frequently than single heterozygotes of either strain, had thin atrophic uteri, had uninterpretable vaginal cytology, and were not able to achieve pregnancy (0 pregnancies out of 7 attempted matings). To test whether suppression of transgene expression with doxycycline could produce healthy and fertile animals, dams were placed on doxycycline chow (40 mg/kg, Bio Serv) prior to mating in an attempt to suppress *in utero* transgene expression in offspring. For pups, doxycycline chow was continued through weaning and sexual maturity (8 weeks of age) and then withdrawn. Although this regimen produced animals with more normal appearing uteri, animals remained infertile and pup mortality on the doxycycline chow was high (> 50%).

Anti-Mullerian hormone receptor 2 (AMHR2) Cre mice$^{+/-}$, which also express Cre in the myometrium, showed a similar phenotype to PR-Cre$^{+/-}$ when crossed with Ai148D$^{+/-}$ (Supplemental Figure 5). We also tried crossing PR-Cre$^{+/-}$ mice with Ai155$^{+/-}$ mice, which express a genetically encoded voltage indicator, QuasAr3, and a channelrhodopsin, CheRiff under dual control of a Cre recombinase and doxycycline (4). These mice expressed the transgenes but also suffered profound adverse animal health consequences.

Finally, we tried electroporation in wild-type mice. Although electroporation of transgenes has been demonstrated in endometrium and smooth muscle (5, 6), we were not able to express transgenes in uterine muscle.

*Virus injection surgery*

Virus was maintained on ice until time of injection. For the virus injection procedure, 8-12 week old heterozygous PR-Cre$^{+/-}$ female mice were deeply anesthetized with 3% isoflurane and then maintained with 1% isoflurane following induction. Body temperature was continuously monitored and maintained at 37 °C with a heating pad (WPI, ATC2000). Eyes were protected from drying with ophthalmic ointment. The abdominal hair was removed with sequential clipping and depilatory cream. Prior to incision, animals were treated with Carprofen (5 mg/kg) and Buprenorphine (0.1 mg/kg). Skin was sterilized with triple application of betadine and alcohol, and a sterile drape applied. Using sterile instruments and a binocular wide-field dissecting microscope, a vertical incision was made left paramedian to the midline to access the peritoneal cavity and the left uterine horn identified. Overlying omental fat and urinary bladder were retracted peripherally using sterile gauze soaked in sterile saline. Additional saline-soaked gauze was applied as needed to prevent drying of the surgical site.

The pattern of trypan dye spread was monitored during each injection to help maximize injection into the muscle layer and minimize luminal or endometrial injection. Repeated injection attempts revealed that dye injected in the muscle layer spread slower (5-10 seconds versus <2 second) and appeared brighter than when injected in deeper layers. When needed, injections were paused and the micropipette repositioned to optimize injection location.

Following injections, Animals were returned to their home cage and treated for 3 days with Carprofen (5 mg/kg) and Buprenorphine (0.1 mg/kg) twice daily. Wound clips were removed on post-operative day 7. Female mice were singly housed for 1-7 days after surgery to protect surgical sites, but always restored to group housing with 1-4 female cage mates by post-operative day 7.

*Live animal imaging*

For live animal imaging, general anesthesia was induced with 3% isoflurane and maintained at 1% isoflurane. Animals were transferred in the supine position to a home-built *in vivo* imaging stage capable of translation and rotation of specimens in three dimensions. Animals were warmed with single-use iron powder hand warmers placed dorsally. Abdominal hair was removed with clippers and depilatory cream. Under a binocular wide-field dissecting microscope, a midline abdominal incision was made to expose the peritoneal cavity and the bladder drained transmurally. Strips of gauze cut from 2×2 inch pads soaked in warmed (37 °C) carbogen (95% $O_2$, 5% $CO_2$)-bubbled KH buffer (Krebs Henseleit: 118 mM NaCl, 4.7 mM KCl, 1.2 mM MgSO4, 1.25 mM $CaCl_2$, 1.2 mM $KH_2PO_4$, 25 mM $NaHCO_3$, 10 mM Glucose bubbled with 95% $O_2$/5% $CO_2$ for pH of 7.4) were used to line the walls of the peritoneal cavity. KH-soaked gauze strips were adjusted to sit between the retractor and the abdominal wall to protect the wall surface from drying or damage from the retractors. Retractors were used to displace bowel superiorly, bladder inferiorly, and uterine mesometerium and associated fat laterally, leaving the uterus exposed centrally. Retractor position was then adjusted as needed to bring the imaged uterine horn fully in view. Strips of KH-soaked gauze were added laterally under the mesometrium and also posteriorly between the uterine horn and any underlying bowel, vessels, or musculature to keep the preparation hydrated and to displace the oviductal end of the uterine horn in the anterior (ventral) direction so as to minimize any posterior displacement of the uterine horn along its longitudinal axis. Care was taken when positioning gauze strips and retractors to avoid putting the uterus under tension. The imaging stage was then moved from the dissecting scope to the functional imaging setup.

*Image processing*

Image analysis was performed in MATLAB. Images were binned in space (2X along each axis), collected as multiple 5-minute-long movies to limit file size, and corrected for camera background. Images were rotated and cropped to produce a consistent orientation with the oviduct of the injected (mouse's left) uterine horn positioned on right. The opposite orientation was used for movies of the uninjected (right) horn.

Image registration was used to correct for changes in jGCaMP8m signal caused by the movement of the organ during contractions. Movies were first segmented into clips (3-166 per video) containing no more than one large contraction episode; episodes identified from changes in mean intensities calculated within regions of interests (ROIs) drawn to segment the organ into thirds. For each clip, a frame near the maximum mean intensity was used to identify feature points which were then tracked sequentially from that frame into each adjacent frame for all frames in the clip using the Kanade-Lucas-Tomasi (KLT) feature-tracking algorithm (7, 8). A geometric transform (local weighted mean) was created from the tracked points for each clip frame, and that transform was then used to register all the frames for each clip.

A polyline was manually drawn along the horn longitudinal axis in each clip, and 10 pixel-long (0.12 mm) by 40-80 pixel-wide (0.49-0.98 mm) regions of interest (ROIs) were tiled along the polyline. Median intensities were calculated for each ROI and each frame, and concatenated for all clips in the movie; where clips differed in the number of ROIs due to changes in organ length, additional ROIs with median intensity values of NaN were added on the oviductal end of the polyline to allow concatenation. The fold change in median fluorescence ($\Delta F/F$) was determined for each ROI relative to the baseline fluorescence, taken as the $10^{th}$ percentile for each ROI, to create kymographs.

For the more active metestrus and diestrus stages, individual calcium event paths were identified from kymographs using Kymobutler, a machine learning software for kymograph analysis (9). Kymobutler was run in Mathematica using the unidirectional model. For the less active proestrus and estrus stages, events paths for each clip were identified from leading half maxima. Propagation distance and velocity, directional bias, locus of initiation, and event rates were determined from event paths. Initiation loci were normalized to the organ length at event onset and inter-event intervals were determined as the time between event initiations, independent of spatial coordinates. Full-width half maxima (FWHM) were determined for each event by interpolating (5-fold) each trace along the event path and averaging to create a per event trace. Power spectra were calculated from each kymograph of $\Delta F/F$ in which the imaged view contained the oviduct. The oviductal signal was defined as the mean value of the fourth, third, and second most rostral spatial bins, and was compared to a representative uterine horn signal defined as the mean value of the central spatial bin and its immediate caudal and rostral neighbor. A fast Fourier transform of these $\Delta F/F$ time sequences was used to generate power spectra of the oviduct and uterine horn for each movie.

*Specimen preparation for whole-uterus explant imaging*

Following live animal imaging, isoflurane anesthesia was continued and an *en bloc* resection of uterus, bilateral oviducts, ovaries, cervix and vagina performed. After specimen removal, the animal was euthanized by isoflurane overdose followed by cervical dislocation. The resected organ was transferred to warmed (37 °C) carbogen-bubbled KH buffer. Any adherent blood was removed with Dumont tweezers, excess mesometrial fat was trimmed with spring scissors, and the specimen was washed three times with fresh KH buffer. The organ was secured through the mesometrium using minutien pins to a dish coated with blackened Sylgard-184 (Dow Chemical, mixed with activated charcoal). The dish was transferred to a heated stage with continuous temperature control (Warner Instruments, TC-344C) and continuously perfused with fresh carbogen-bubbled KH at 4 mL/min, maintained at 37 °C by an inline heater (Warner Instruments, SF-27-B).

Except where noted, whole explant imaging was always performed following live animal imaging. 20-40 minutes was typically required between ending a live animal imaging session and starting an explant imaging session. Explant imaging sessions varied in length between 30-120 minutes.

*Calcium imaging of whole uterine explants*

Explanted preparations have been widely used to study uterine physiology, but their activity may not faithfully mimic *in vivo* activity. The uterus comprises an inner layer of circumferential muscle fibers and an outer layer of axially elongated fibers. Thus slicing along any axis will cut some of these fibers, possibly perturbing activity patterns. To avoid cutting across myometrial fibers, whole uterine explants were removed *en bloc*, comprising vagina, cervix, uterus, oviducts, and ovaries. These preparations were washed, warmed, and perfused with oxygenated solutions and imaged using the same approach and microscope as used for *in vivo* imaging (Methods). Explanted uteri were largely quiescent regardless of cycle stage, a marked difference from the *in vivo* recordings (Supplemental Figure 4). Transient activity could be induced by mechanical stimulation of the explant (Supplemental Figure 4) or electrical shock, but this activity also was not similar to what was observed in live animals. In an attempt to recreate *in vivo*-like activity patterns, we let the explants equilibrate for up to 2 hours in warmed and oxygenated solution. We also tried explanting organs immediately after induction of anesthesia, rather than following an *in vivo* imaging session. Both approaches yielded quiescent explant preparations, markedly different from the live animal preparation. From these experiments, we conclude that there may be important circulating factors or mechanical stimuli present *in vivo* which are absent from standard *ex vivo* live-tissue preparations.

*Preparation of uterine rings for immunofluorescence and confocal microscopy*

Following live-animal imaging, isoflurane anesthesia was continued and an *en bloc* resection of uterus, bilateral oviducts, ovaries, cervix and vagina performed. The resected organ was transferred to an ice-cold solution of PBS, cleaned of any adherent blood with Dumont tweezers under a dissecting microscope, and then transferred to a 4% paraformaldehyde solution in PBS for 30 minutes. The animal was euthanized. Fixed uteri were then washed three times in ice cold PBS, transferred to a plastic 35 mm culture dish, and immobilized in 4% agarose matrix. Agarose blocks containing segments from the injected and contralateral horn were cut and then sliced into 100 micron rings using a vibratome (Leica VT1000).

### Supplemental Discussion

No transgenic Cre lines have been reported which drive expression solely in the myometrium, and there are no reported plasmid-based specific myometrial promoters. The adverse effects seen here with both PR-Cre and AMHR2-Cre driver lines tested could reflect effects of reporter overexpression in non-uterine tissues (*e.g.* ovaries), the TTA promoter system in both tested reporter strains, or chronic and high-level overexpression of GCaMP6f. Chronic GCaMP6 expression in the brain of transgenic mice can cause seizure-like activity patterns (10), Myometrial transgene expression or transcript silencing has been previously accomplished with adenoviral and lentiviral vectors (11, 12). However, compared to AAVs, adenovirus produces weaker and less durable construct expression, as well as host inflammatory responses while lentivirus carries risks of insertional mutagenesis (13).
